# Acid cleavable biotin-alkyne improves sensitivity for direct detection of biotin labeled peptides in BONCAT analysis

**DOI:** 10.1101/2024.07.16.603801

**Authors:** Daniel B. McClatchy, John R. Yates

## Abstract

BONCAT (Biorthogonal noncanonical amino acid tagging) is a labeling strategy that covalently adds a biotin-alkyne (BA) to methionine analogs via a click reaction. When methionine analogs are incorporated into a proteome, enrichment of the BA-labeled proteins allows the detection of newly synthesized proteins (NSP) by mass spectrometry. We previously reported that using our Direct Detection of Biotin-containing Tags (DidBIT) strategy, protein identifications and confidence are increased by enriching for BA-peptides instead of BA-proteins. We compared cleavable BA (DADPS) and uncleavable BA in the identification and TMT quantification of NSP. More than fifty percent more proteins were identified and quantified using DADPS than with uncleavable BA. Interrogation of the data revealed that multiple factors are responsible for the superior performance of DADPS.

## 1. Introduction

Mass spectrometry (MS) based proteomic analysis of complex biological samples has led to numerous exciting biological discoveries by revealing changes in protein expression induced by biological perturbations. These advances stem from mass spectrometry’s ability to identify and quantify proteins involved in novel biological processes in a global and unbiased manner. As biologists perform more in-depth investigations of the molecular underpinnings of cell physiology, they face the challenge of identifying smaller and more subtle changes in complex proteomes such as tissue.

Biorthogonal noncanonical amino acid tagging (BONCAT)^1^ is a protein labeling strategy that has been employed to address this challenge. BONCAT employs non-canonical amino acids (ncAA) so that only proteins synthesized within a discrete temporal window willbe analyzed by LC-MS^1^. It assumes that newly synthesized proteins (NSP) are the first sub-proteome to respond to biological perturbations and that the presence of older proteins obscures the detectable changes in the newly synthesized proteome when the “old” and newly synthesized proteins are analyzed together, as in a typical proteomic experiment. Thus, BONCAT can detect changes that may be obscured in whole proteome analysis. Azidohomoalanine (AHA), was first reported in 2006 and has been the most widely used ncAA since then. AHA is inserted into proteins through interaction with the endogenous methionine (Met) tRNA synthetase (MetRS). Because Met competes with AHA, Met depletion greatly increases AHA incorporation into the proteome^1^. However, since Met is an essential AA, extended AHA labeling times without Met could be deleterious^2^. Although initially used in cultured cells, AHA has now been used to label more complex proteomes, such as C.elegans and mice^3, 4^. AHA labeled proteins have identical structure^5^, stability^6^, and cellular localization^7^ as native unlabeled proteins, and the function of AHA labeled proteins has been shown to be indistinguishable from endogenous unlabeled proteins. Thus, the functional AHA proteome and the unlabeled proteome in cultured cells^8^, primary neurons^9^, liver^4^, heart^10^, and embryonic murine development^11^ respond identically to perturbations. Novel changes identified with AHA proteins have been verified with unlabeled endogenous proteins ^12–14^. The use of azidonorleucine (ANL) as a ncAA was first reported in 2016^15^. Even though ANL is Met analog, it requires a mutated MetRS (mMetRS) for incorporation into the proteome. There have been conflicting reports about whether endogenous Met interferes with ANL incorporation ^15–17^. Since only cells with mMetRS will incorporate ANL, specific cell types can be chosen for proteomic analysis. Alvarez-Castelao et al. took advantage of this selectivity to create mice with Cre-recombinase induced expression of mMetRS in specific cell types in the brain^18^. Although the quantitative analysis of specific cell types in tissues has the potential to drive numerous biological discoveries, it will require more sensitive protocols than those previously used for AHA.

In our BONCAT protocol, we employ the peptide enrichment strategy DidBIT (Direct Detection of Biotin-containing Tags)^19^. In DidBIT, the AHA-BA peptides are enriched with neutravidin beads. The eluted AHA-BA peptides are then analyzed by LC-MS. Others have confirmed that peptide enrichment is superior to protein enrichment (i.e. neutravidin enrichment prior to digestion) for biotin tags^20, 21^. Since the interaction between biotin and neutravidin is extremely strong, the optimization of the AHA-BA peptide elution from the neutravidin beads has the potential to increase the sensitivity of the DidBIT strategy. In this study, we compared the sensitivity of a cleavable and uncleavable BA in our DidBIT-BONCAT protocol.

## 2. Methods and Materials

### Cell Culture

Neuro2A (Figure 1A-C) and HEK293 (Figure 1 D&E) cells were maintained at 37°C with 5% CO_2_. HEK293 cells were cultured in DMEM with GlutaMAX (Gibco # 10569-10) with 10% FBS added. Neuro2A cells were cultured in EMEM with L-Glutamine (ATCC# 30-2003) with 10% FBS added. mMetRS-Flag-mCherry (a gift from David Tirrell (Addgene plasmid # 89189; http://n2t.net/addgene:89189; RRID:Addgene_89189)) was transiently transfected into the cells using XtremeGene HP transfection reagent(Roche). Forty-eight hours after transfection, the cells were harvested for analysis. To label with ANL(Tocris) or AHA(Click Chemistry Tools), the previous published protocol was followed^22^. Briefly, cells were incubated in Hank’s Balanced Salt Solution (HBSS) with Glutamax, sodium pyruvate, 4mM MgCl_2_, 4mM CaCl_2_ for 30minutes, then cells were incubated with 1mM of ANL or AHA for 1hour in HBSS. For experiments testing the effect of methionine, 15μg/ml of methionine was added to the HBSS. Cells were harvested by scraping them off the plate in PBS and were then lysed in PBS with a tip sonicator (Qsonica).

**Figure 1.**
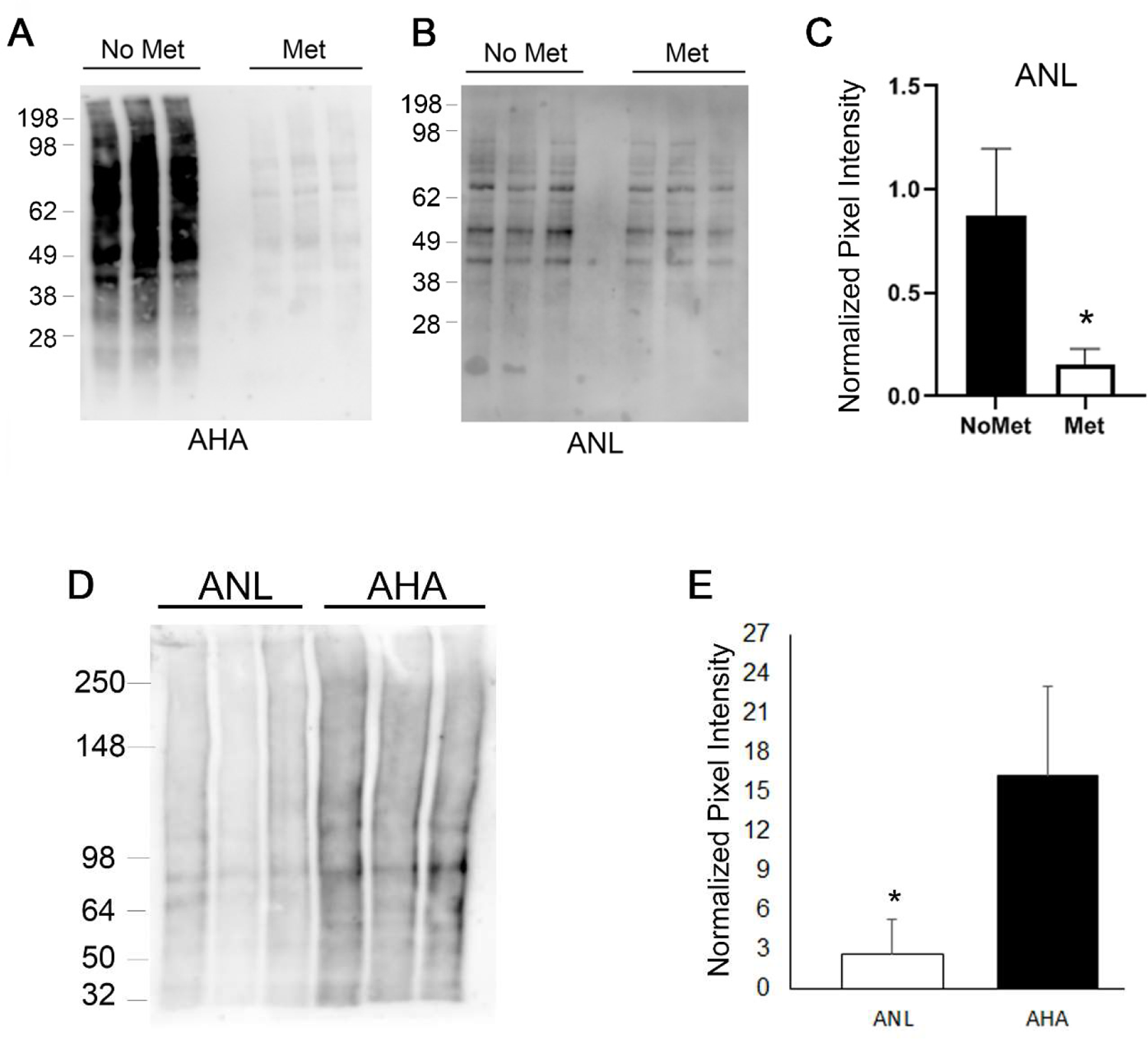
Comparison of ANL and AHA incorporation in cultured cells. **A**, AHA incorporation is abolished when methionine is present. **B**, The presence of methionine only slightly reduced ANL incorporation. **C**, Quantification of ANL samples shown in Fig1A. **D,** Immunoblot analysis of FACS. **E,** Quantification of samples shown in Fig1D. Each lane on the immunoblots represents a different biological replicate and immunoblots were probed with streptavidin-HRP. * p< 0.05.

### Immunoblot Analysis

*After addition of 4X Laemmli Sample Buffer with β-mercaptoethanol, click reactions were separated on a gradient gel (4-12% Bis-Tris or 3-8% tris-acetate)were transferred to PVDF blotting paper.* The immunoblotting antibodies were β-actin (Sigma#A5441), FLAG-tag(VWR# 95040-926), and Streptavidin-HRP(JacksonImmuno) and rabbit anti-biotin antibody(Bethyl laboratories). The immunoblots were visualized with Azure Biosystems Radiance ECL reagent and Azure Biosystems c600 imaging system and were quantitated as previously described^23^. To determine the total protein content of the samples, gels were stained with AzureRed Fluorescent Protein Stain (Azure Biosystems). For quantification, immunoblot signals were normalized to the protein stain before statistical analysis.

#### FACS

HEK293 cells transfected with mMetRS-mcherry were isolated using a Beckman-Coulter Astrios cell sorter with 488nm excitation for scatter and 565nm excitation to detect mCherry emissions using a 614/20nm bandpass filter. To accommodate the larger cell size, the sorter was configured to use a 100μm nozzle size and a lower sort pressure of 25psi. FACS was performed in the Scripps Research core.

### Animals

Mice were housed in plastic cages located inside a temperature- and humidity-controlled animal colony. Animal facilities were AAALAC (Association for Assessment and Accreditation of Laboratory Animal Care) approved, and protocols were in accordance with the IACUC(Institutional Animal Care and Use Committee). AHA labeled mice were male C57BL/6 2month old mice from the TSRI colony. For ANL tissue labeling, C57BL/6-Gt(ROSA)26Sortm1(CAG-GFP,-Mars*L274G)Esm/J mice were bred with B6.Cg-Tg(Camk2a-cre)T29-1Stl/J to obtain neuronal specific expression. The initial breeder mice to start the colony were obtained from Jackson laboratories. The mice were labeled with AHA or ANL via the diet as previously described^4^. The mice were anesthetized with halothane and sacrificed by decapitation. The brains were quickly removed and dissected, then snap-frozen with liquid nitrogen. Brains were homogenized in a Teflon dounce grinder using PBS with proteases inhibitors without EDTA.

### DidBit-BONCAT

Prior to the click reactions, Pierce BCA Protein assay was performed on all samples. The click reaction was performed as previously described except for minor modifications as follows^4^. Biotin-PEG4-alkyne (i.e. UnC) and DADPS were purchased from Click Chemistry Tools. The click reaction was incubated at 30°C for 1hour, then incubated at 4 °C overnight while rotating. The click reactions were then precipitated with cold acetone overnight at −20°C. The dried precipitated protein pellets were resuspended with 8M urea in 100mM HEPES, pH 8.5. The proteins were then reduced with 5mM TCEP for 20min at 55 °C, then alkylated with 10mM chloro-iodoacetamide for 20min in the dark at room temperature. The urea was then reduced to 2M with 100mM HEPES, pH 8.5. Trypzean (Sigma) was added at 1:25 ratio to protein amount, then incubated overnight on a 37 °C shaker. The digestions were centrifuged for 15min at 21,000 x g and the supernant was incubated with 50ul of neutravidin beads overnight at 4 °C. The beads were washed three times with PBS followed by twice with deionized water.

DADPS click reactions were eluted with 5% formic acid for 30min while shaking at room temperature. This elution was performed twice, and the eluates were combined into one tube. UnC click reactions were eluted with 80% acetonitrile, 0.2% TFA, and 0.1% formic acid as previously described^22^. The UnC elution was dried with a speed-vac, then UnC dried peptides and DADPS elution were desalted with the Sep-Pak tC18 elution plate (Waters). For TMT labeling, dried peptides were resuspended in 8ul of 0.5M HEPES, pH 8.5. Each TMT label was resuspended in anhydrous acetonitrile for a concentration of 10ug/ul. One microliter of TMT was added to resuspended peptides and incubated for 30 min, then this step was repeated. The labels were quenched with 5% hydroxylamine. The TMT labeled peptides were desalted prior to LC-MS analysis.

### BA-peptide enrichment with a Biotin Antibody

The enrichment of biotinylated peptides by antibodies was performed as previously described^21^. Briefly, dried desalted peptides were resuspended in 50 mM MOPS (pH 7.2), 10 mM sodium phosphate and 50 mM NaCl. BA-peptides were enriched with 50μg of Biotin Antibody Agarose (ImmuneChem Pharmaceuticals Inc.) for 1 hour at 4°C. After the agarose was washed, BA-peptides were eluted with 0.15% TFA for 5min at room temperature. All reagents and samples were kept on ice during the protocol.

### LC-MS Analysis

Desalted dried peptides were resuspended in 30ul buffer A (5% acetonitrile, 95% UltraPure H_2_O and 0.1% formic acid), and bath sonicated for 30min. Ten microliters of each sample was injected directly onto a 25 cm, 100 µm ID column packed with BEH 1.7 µm C18 resin (Waters) after equilibration with 20 µl of buffer A. Samples were separated at a flow rate of 400 nl/min on a nLC 1000 (ThermoFisher) using a gradient of 1-25% buffer B over 120 min, an increase to 40% buffer B over 40 min, an increase to 90% buffer B over 10 min and holding at 90% buffer B for a final 10 min, for total run time of 180 min. Buffer A and B were 0.1% formic acid in 5% and 80% acetonitrile, respectively. Peptides were eluted directly from the tip of the column and nano-sprayed directly into the Fusion Eclipse or Lumos Orbitrap tribrid mass spectrometer (ThermoFisher) by application of 2.5 kV voltage at the back of the column. The mass spectrometer was operated in a data-dependent mode. Full MS scans were collected in the Orbitrap at 120K resolution with a mass range of 400 to 1500 m/z and an AGC target of 4e5. The cycle time was set to 3s, and within this 3s the most abundant ions per scan were selected for CID MS/MS in the ion trap with an AGC target of 2e4 and minimum intensity of 5000. Maximum fill times were set to 50 ms and 100 ms for MS and MS/MS scans, respectively. Quadrupole isolation at 1.6 m/z was used, monoisotopic precursor selection was enabled, and dynamic exclusion was used with exclusion duration of 5 sec. For the TMT 10plex samples, MS3 analysis with multinotch isolation (SPS3) was utilized for detection of TMT reporter ions at 60k resolution on the Eclipse.

### Data analysis of MS data

MS/MS spectra were searched with the Prolucid algorithm^24^ against the Uniprot mouse database (Release date 04-24-2018) that was concatenated to a decoy database in which the sequence for each entry in the original database was reversed. A mass shift of 57.02146 on cysteine wass used as a static modification in searches. Differential modification of the methionine residue as the result of AHA-BA mass shift was set at 79.0711 for DADPS or 452.2376 for UnC. Searches allowed for partial tryptic status and two miscleavages. The search results were assembled and filtered using the DTASelect 2.0 program^25^. For TMT 10plex experiments, Proteome Discoverer 2.5 was used for identification and quantification. For TMT experiments, the TMT mass shift of 229.1629 was searched as a static modification for lysines and the peptide N-terminus, and an AHA-BA mass shift was searched as a differential modification. Peptides were required to be partially tryptic and less than 10ppm deviation from peptide match. A global FDR at the protein level of 0.01 was applied. PACOM was used for determining unique protein and peptides, as shown in Figure 2B^26^. Graphs and statistical analyses were performed in Prism Graphpad and Microsoft Excel.

**Figure 2.**
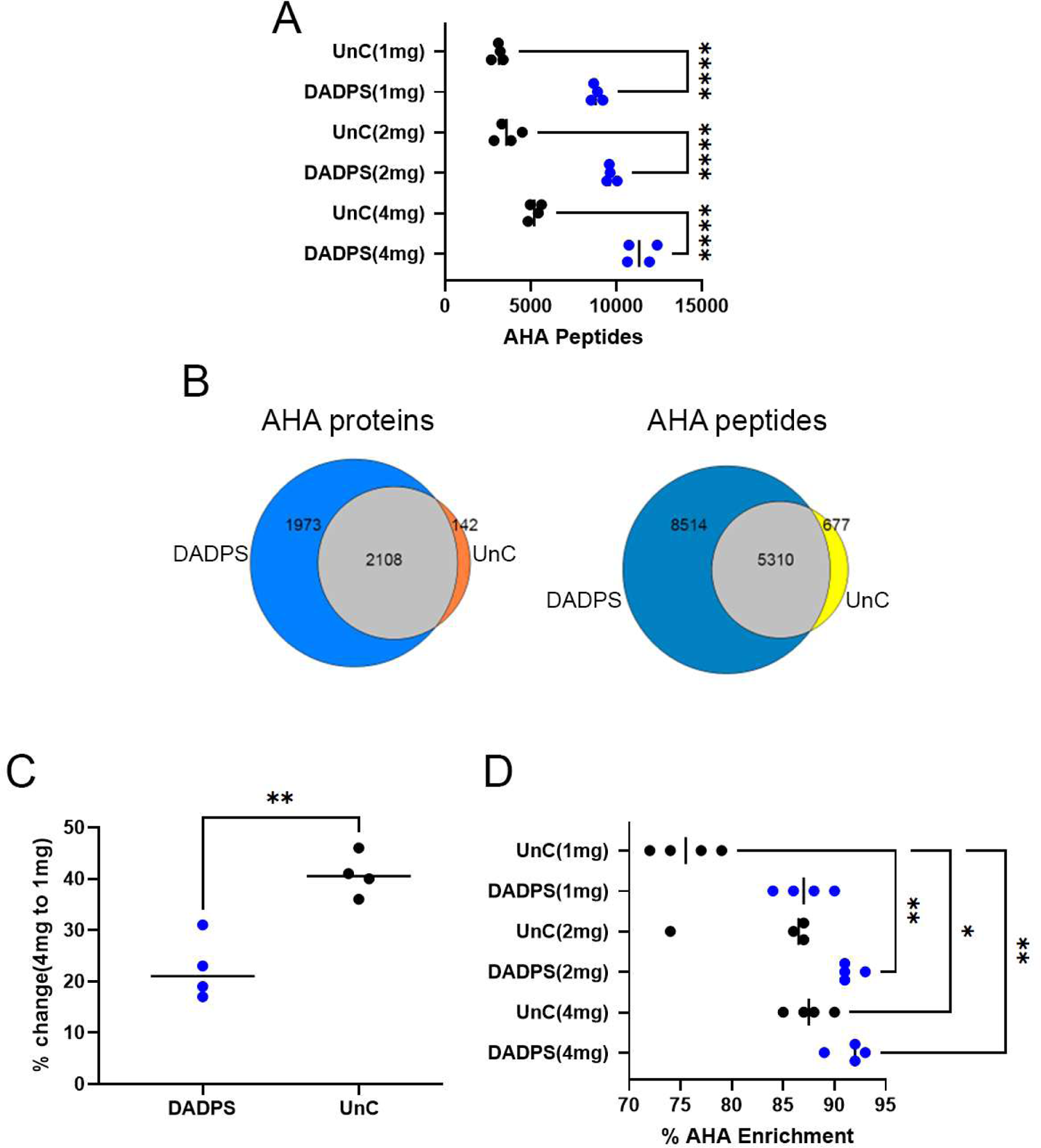
**A,** Identifications of AHA-BA modified peptides using either DADPS or UnC BA. **B,** The overlap of unique AHA protein and peptide identifications between the DADPS and UnC. **C**, Percent decrease in AHA peptide identifications from 4 to 1mg samples using DADPS or UnC. **D**, Percent enrichment of AHA peptides from the analysis. * p< 0.05, ** p< 0.01, *** p < 0.001, **** p < 0.0001.

## 3. RESULTS

### ANL incorporation in the proteome is less efficient than AHA in cultured cells

ANL and AHA labeling were compared in cell culture. In the first experiment, mMetRS tagged with mCherry and FLAG^15^ was transfected into cultured cells, then the cells were labeled for 1 hour with 1mM ANL or 1mM AHA in media with or without Met. Click reactions were then performed to conjugate the BA onto proteins containing AHA or ANL. Immunoblot analysis was performed with streptavidin-HRP to determine the extent of the ncAA incorporation. As previously reported, AHA incorporation was almost completely prevented when Met was in the labeling media, and it could not be quantitated without saturation of the signal of the No-Met samples (Figure 1A; Supplementary Figure 1A). In contrast, the addition of Met produced a small but significant (p=0.019) decrease in ANL incorporation (Figure1B, C and Supplementary Figure 1B). Since ANL incorporation is limited by the transfection efficiency of mMetRS and AHA incorporation is not, this experiment is not a fair comparison of the incorporation rates. Next, cells were transfected with mMetRS and then labeled with 1mM ANL or 1mM AHA for 1hour without Met. Transfected cells were enriched by Fluorescence-activated Cell sorting(FACS) using the mCherry tag. FACS increased the detection of mMetRS and ANL (Supplementary Figure 1C, D,E). Since AHA does not rely on mMetRS for incorporation, FACS had no effect on AHA incorporation (Supplementary Figure 1D,E). However, the AHA protein signal was almost five times greater than ANL on immunoblots (Figure 1D, E and Figure S1E). This suggests that AHA incorporates into the proteome more efficiently than ANL. The most plausible explanation is that endogenous MetRS charges the Met tRNA more efficiently than the exogenous expressed tagged mMetRS. Therefore, detection of ANL will require a more sensitive protocol than AHA because it is only incorporated into the proteomes of a specific cell type and it is incorporated less efficiently..

### Cleavable BA outperforms uncleavable BA

To develop a more sensitive DidBIT-BONCAT protocol, we compared the acid cleavable biotin-alkyne dialkoxydiphenylsilane (DADPS, C_42_H_62_N_4_O_9_SSi) and the uncleavable (UnC) biotin-alkyne Biotin-PEG-alkyne (C_12_H_35_N_3_O_6_S). UnC has been successfully employed in published BONCAT LC-MS studies^4, 10, 27^. Three different quantities of AHA labeled brain tissue (4, 2, and 1mg) were analyzed. Four replicates of each of the 4, 2 and 1mg samples were processed independently with either DADPS or UnC for a total of 24 samples. After the click reactions, samples were digested and then enriched with neutravidin beads. DADPS was eluted with 5% formic acid and UnC was eluted with 80% acetonitrile, 0.2% trifluoroacetic acid, and 0.1% formic acid. After elution, each sample was analyzed once on an Orbitrap Eclipse mass spectrometer after a 3hour reversed-phase gradient. The differential modification on Met was searched to identify the AHA-DADPS (i.e. 79.0711) or AHA-UnC (i.e. 452.2376) modified peptides. We identified >50% more AHA-BA peptides with DADPS than with UnC for all starting amounts (Figure 2A). After filtering for unique AHA peptides, the majority (89%) of the UnC AHA peptides were identified in the DADPS dataset (Figure 2B). As expected, we observed a decrease in AHA peptide identifications with less starting material for both BA. However, UnC was more sensitive to the amount of starting material. UnC identified 41% less AHA peptides in 1mg samples than in 4mg samples, whereas DADPS identifications only decreased by 22% (Figure 2C). We also analyzed the percentage enrichment of AHA, which was defined as the total number of AHA peptides identified divided by sum of AHA and unmodified peptides identified. For 2mg and 4mg samples, DADPS and Unc enrichment was comparable, but with 1mg of starting material the average percentage enriched for UnC-BA was 76% and for DADPS it was 87% (Figure 2D). The reproducibility of AHA-DADPS peptide identifications was slightly better than UnC (Supplementary Figure 2A). DADPS was also superior to UnC when we used 1.5mg of brain tissue labeled with ANL (Supplementary Figure 2B).

### TMT labeling combined with DidBIT-BONCAT

We examined the compatibility of the BA with TMT isobaric labeling. First, we tested whether TMT labeling affects the AHA-BA peptide identification by repeating the previous experiment on peptides that were labeled with TMT after elution from the neutravidin beads. AHA labeled brain tissue samples (i.e., 4, 2, and 1mg) were independently processed three times with either DADPS or UnC for a total of 18 samples. Using the sample MS acquisition parameters, we identified fewer AHA peptides than in the previous non-TMT AHA experiment for both DADPS and UnC. It has previously been reported that TMT labeling can reduce identifications^28, 29^. We also observed >60% more AHA peptides with DADPS than with UnC (Figure 3A). Next, we performed 10plex TMT experiments, using 0.2 mg of starting material for each TMT channel. Each channel was processed separately and the AHA peptides were labeled with TMT after the neutravidin elution. For each BA, we performed three 10plex experiments and employed the SPS-MS3 method^30^ for data acquisition. DADPS identified 2.8 times more peptides and quantified 4.4 times more peptides than UnC (Figure 3B). As shown in Figure 2D AHA peptide enrichment efficiency was lower in the TMT experiment than in the previous experiment, consistent with our observation that lower amounts of starting material (i.e. 0.2mg) decrease the efficiency of the AHA peptide enrichment. However, AHA enrichment efficiency for the DADPS samples was still significantly higher than for UnC samples(Figure 3C). We examined the PSMs to determine which quantification filters (i.e. signal to noise or SPS MS3 mass matches) were responsible for the poor quantification of UnC AHA peptides. For a PSM to be considered for quantification, a signal to noise (S/N) of 10 was required and at least 65% of SPS MS3 matches. We did not observe a difference of the S/N distribution between DADPS and UnC PSMs (Figure 3D). Examining the percentage of SPS MS3 mass matches to the identified peptide, we observed that DADPS had more PSMs that possess more 65% or greater matches than UnC.

**Figure 3.**
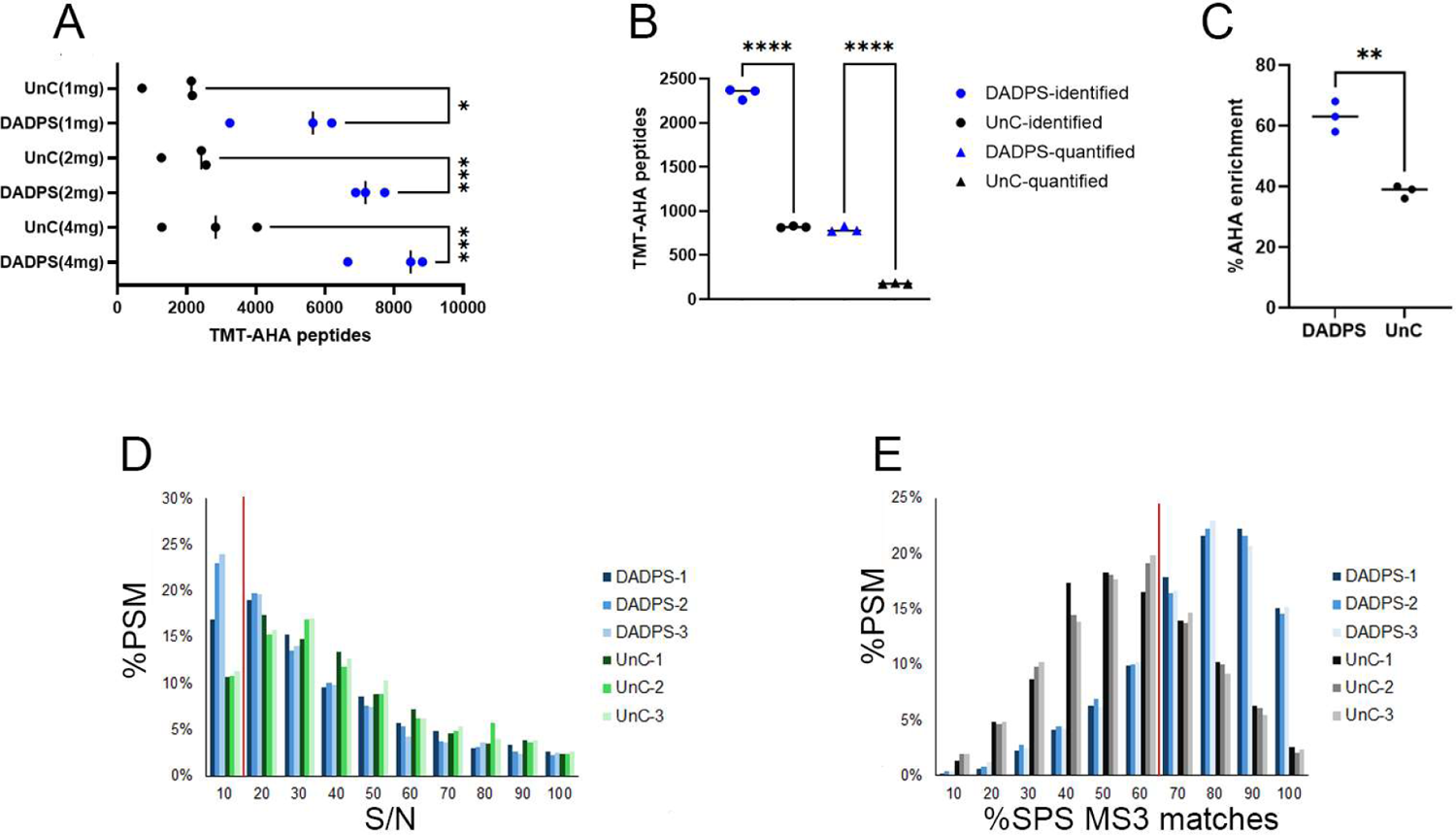
**A,** Identification of TMT-AHA peptides with a single TMT label using MS2 acquisition **B,** Identification and quantification of TMT-AHA peptides using 10plex TMT labels with SPS MS3 acquisition. **C,** The percent AHA enrichment of the identified peptides in B. **D**, The S/N of the TMT-AHA PSMs acquired. Each replicate is plotted individually. The red line indicates the required S/N (i.e. 10) threshold for quantification. **E**, The percentage of SPS MS3 matches of the TMT-AHA PSMs. Each replicate is plotted individually. The red line indicates the required SPS MS matches (i.e. 65%) threshold for quantification. * p< 0.05, ** p< 0.01, *** p < 0.001, **** p < 0.0001.

### AHA-UnC peptides have different properties than AHA-DADPS peptides

We sought to determine the reason for the superior performance of DADPS. When we compared the protein lysates after click reactions of DADPS vs UnC on immunoblots, we did not observe an increase in biotin signal with DADPS(Figure 4A). Peptides from 2mg of AHA labeled brain were quantitated after the neutravidin elution using a BCA peptide assay. DADPS was calculated to have average of 71.7ug/ml peptides per elution whereas UnC-BA had 62.5 ug/ml(Figure 4B), but this difference was not significant. Although this difference suggests that DADPS elution is more efficient than UnC-BA elution, it cannot account for the difference in peptide identifications (>2x).

**Figure 4.**
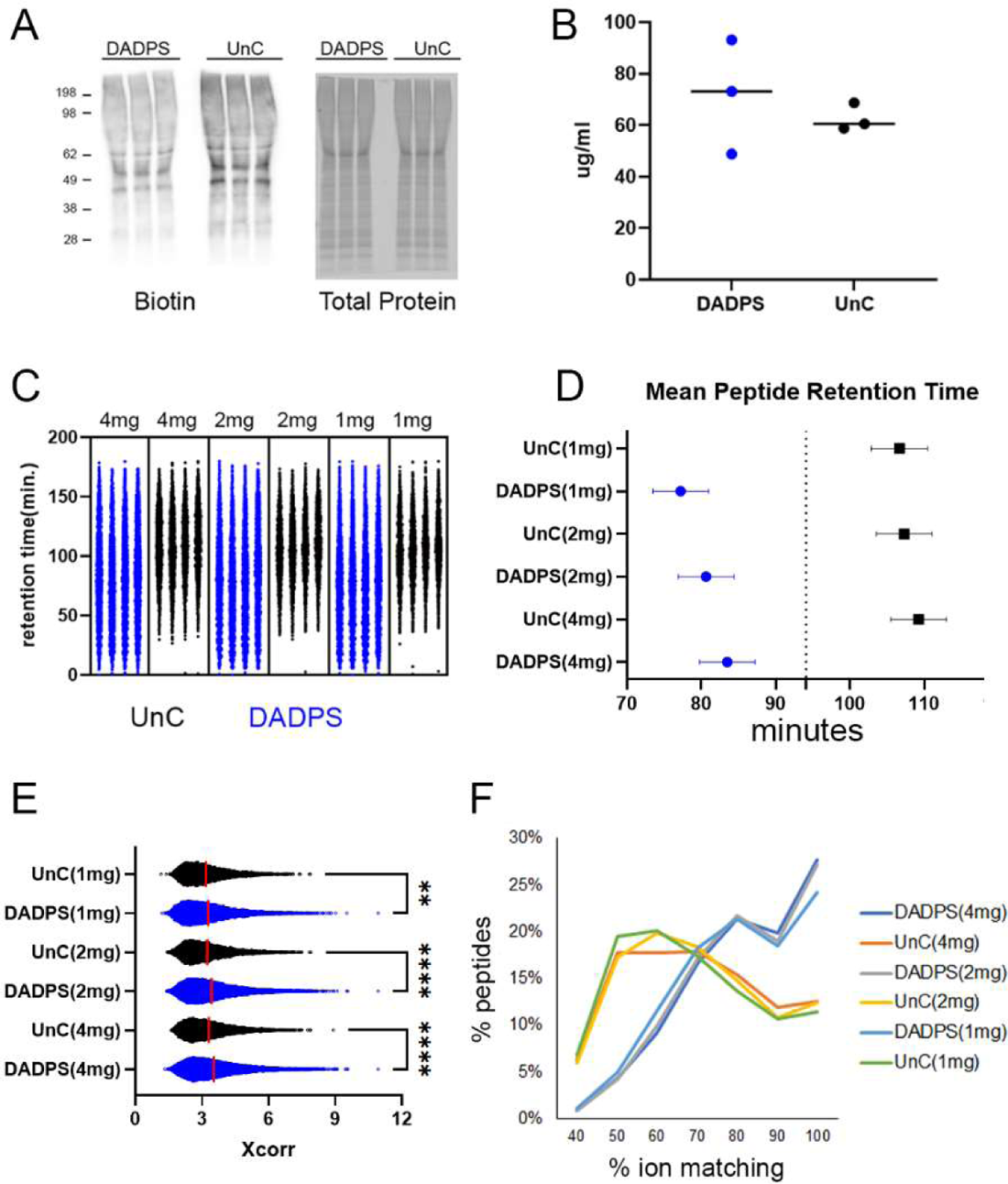
**A,** Immunoblot of click reactions with either DADPS or UnC using a biotin antibody. The same samples were analyzed on a separate gel and stain with a total protein stain. **B**, Peptides eluted from neutravidin beads were quantitated with a peptide BCA assay. **C**, Retention time (RT) distribution of AHA-BA peptides. DADPS (blue), UnC(black). **D**, Median RT of AHA-BA peptides. **E**, Distribution of Xcorr scores assigned to identified AHA peptide modified with either DADPS or UnC. **F**, Percentage of ions matching theoretical MS2 spectra from identified DADPS or UnC peptides. * p< 0.05, ** p< 0.01, *** p < 0.001, **** p < 0.0001.

Next, we investigated the MS data for differences in the DADPS and UnC-BA peptides. We observed a striking difference in the chromatographic profiles of the DADPS and UnC peptides. DADPS peptides were evenly distributed in the chromatogram whereas UnC-BA peptides appeared later in the chromatogram (Figure 4C). Specifically, compared to DADPS peptides, the retention times (RT) for UnC peptides were delayed more than 20 minutes (Figure 4D). DADPS peptides were assigned significantly higher Xcorr scores than UnC peptides (Figure 4E). Correspondingly, the proportion of ions in the MS2 spectra that matched the theoretical spectra was reduced in the UnC peptides compared to the DADPS peptides (Figure 4F). The lower Xcorr scores and matching ions suggest that the UnC peptides are not being as efficiently fragmented as the DADPS peptides. We attempted to improve the fragmentation of the AHA-UnC peptides by using HCD instead CID, but we did not observe an increase in peptide identifications (Supplementary Figure 2C). Charge state distributions also differed between the BA; 54% of the DADPS PSM were +2 whereas 39% of the UnC-BA PSMs were +2 (Figure 5A). We investigated whether the altered charge state distributions of the UnC peptides were related to its other characteristics of delayed RT, lower Xcorr scores, and lower ion matching. When the RT was analyzed for different charge states, +3 and higher charged peptides had significantly delayed RT compared to +2 peptides for both DADPS and UnC peptides (Figure 5B). But when peptides of the same charge state were compared, UnC peptides still had significantly delayed RT compared to DADPS peptides. Similarly, when comparing peptides of the same charge state, Xcorr scores and percentage ion matching were significantly lower with the UnC peptides (Figure 5C and D). Thus, a larger percentage of +3 and higher charged peptides in the UnC data does not appear to be related to its other distinguishing biochemical characteristics.

**Figure 5.**
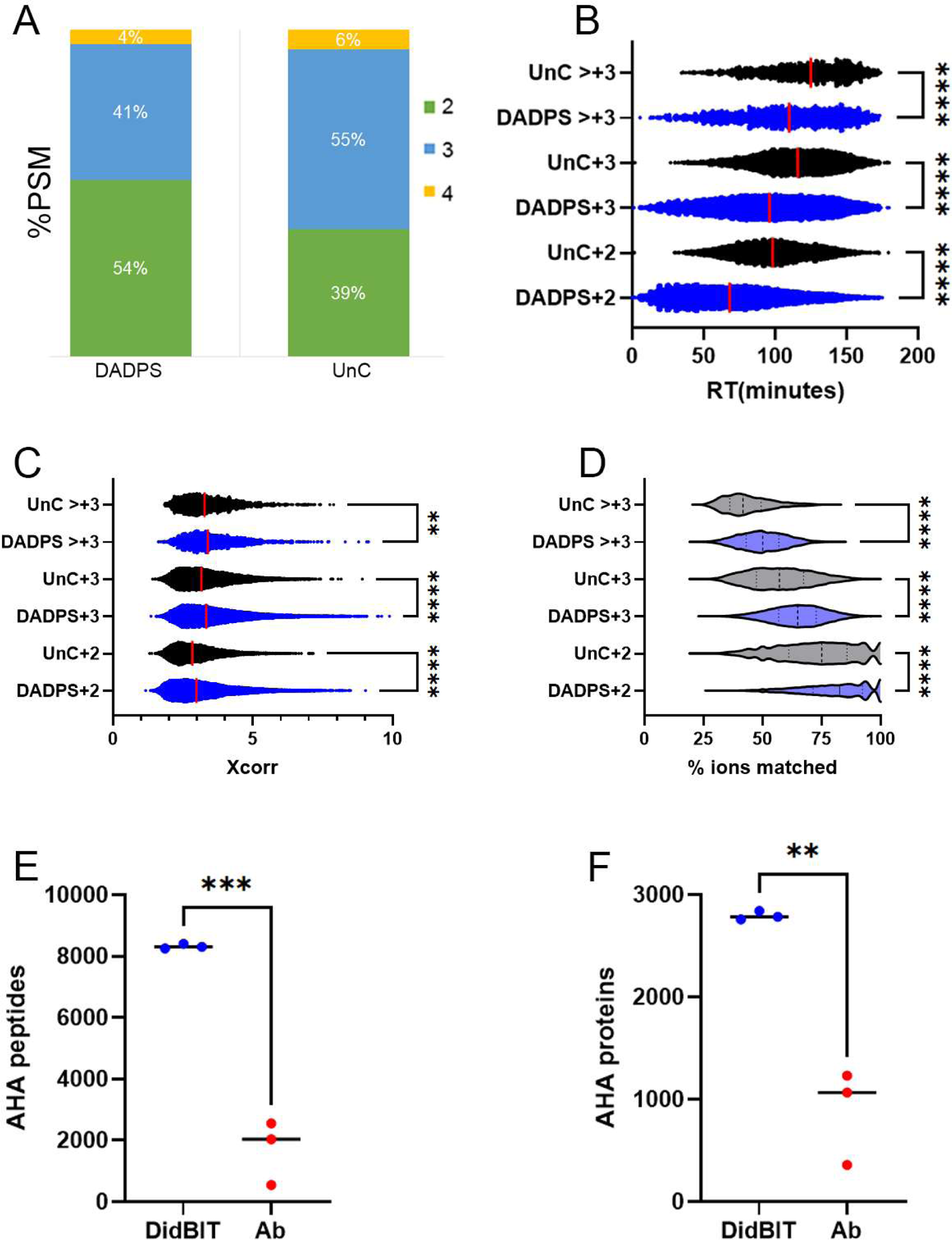
**A,** Percentage of PSM with their assigned charge states for the DADPS and UnC data in Figure 1A. **B,** Retention times (RT) for DADPS and UnC peptides separated by peptide charge states. **C,** Xcorr scores for DADPS and UnC peptides separated by peptide charge states. **D**, Violin plot of the percent ions matching for DADPS and UnC peptides separated by peptide charge states. The number of AHA peptides (**E**) and AHA proteins (**F**) identifications with DidBIT and Ab enrichment strategies.

### Comparison of neutravidin enrichment with antibody (Ab) enrichment

We compared our DidBIT-BONCAT protocol with DADPS with Ab enrichment. Ab enrichment of biotinylated peptides has been reported to identify more modified peptides than neutravidin enrichment^20, 21^. It was postulated that the lower binding affinity of a biotin vs neutravidin Ab for biotinylated peptides is the basis for improved performance, but these comparisons were not performed using a BONCAT strategy (i.e. biotin-alkynes). Since the Ab strategy employs acid to elute biotinylated peptides from biotin Ab attached to beads, DADPS is not compatible with Ab enrichment. Therefore, we compared DADPS-AHA peptide neutravidin enrichment to UnC-AHA peptide Ab enrichment^21^. For each strategy, one milligram of AHA labeled brain tissue was processed independently three times. DidBIT identified almost 80% more AHA peptides than the Ab enrichment strategy and 68% more AHA proteins (Figure 4E and F).

## 4. Discussion

The binding between biotin and avidin is one of the strongest in nature^31^, which is why biotin is a popular choice in many enrichment strategies in biological research.

However, elution of biotin moieties from avidin solid supports is challenging. We suspect that biotinylated peptides may elute more readily than biotinylated proteins, which may be one reason that the DidBIT peptide enrichment strategies outperforms biotin protein enrichment strategies^19^. In this study, we hypothesized that a cleavable BA would improve the sensitivity of the DidBIT protocol. DADPS was chosen because it has been shown to be superior to other cleavable biotin reagents for LC-MS analysis^32, 33^. To our knowledge, DADPS has not been used in any BONCAT studies. We observed a striking increase in the identification and quantification of BA-peptides with DADPS. Two laboratories have directly compared the efficiency of neutravidin (i.e. DidBIT) and a biotin specific Ab for biotinylated peptide enrichment and both concluded that the biotin Ab is more efficient^20, 21^. These reports examined biotin reagents that did not include biotin-alkynes. Since the biotin Ab enrichment strategy elutes peptides with acid, it is not compatible with DADPS. The large biotin fragment (> 700Da) of DADPS would be eluted into the mass spectrometer with the AHA peptides, whereas the biotin moiety is retained on the neutravidin agarose using DidBiT. Direct comparison of DidBIT with DADPS and biotin Ab with UnC demonstrated an 80% increase in AHA peptide identifications with DidBIT. Thus, we conclude that the DidBIT strategy employing DADPS is the best choice for identifying BA peptides in BONCAT experiments.

We assumed the improved performance of DADPS stems from a more efficient elution from the neutravidin beads. While we observed a more efficient elution, the modest increase over UnC was not enough to explain the large increase in AHA peptide identifications. So, we investigated further to identify other reasons for the superiority of DADPS. We observed more than a 20-minute delay in RT for the UnC peptides compared to DADPS, which comports with previous report that biotinylated peptides have delayed RT compared to unmodified peptides^34^. This RT delay is most likely due to the increased hydrophobicity caused by the biotin moiety and suggests that the detection of UnC peptides could be improved by altering the LC gradient for better separation of hydrophobic peptides. Specifically, shallow gradients at higher organic solvents should be investigated. However, optimization of LC gradients would probably not increase the number of detected UnC peptides to DADPS levels because UnC peptides are influenced by other properties that hinder their identification. The Xcorr scores and ion matching of UnC spectra were lower for UnC spectra than for DADPS spectra, suggesting poor fragmentation of the UnC peptides. This may stem from the large UnC tag, which is five times larger than the DADPS tag. It is possible that optimizing the collision energy could improve UnC fragmentation and lead to more peptide identifications. Finally, we observed a decrease of +2 charged peptides and increase of +3 and higher charged peptides in the UnC MS data. This may decrease identifications since database searching algorithms performed better on doubly charge peptides^35^. Alternatively, it has been reported that CID fragmentation favors +2 charged peptides^36^. When we compared +3 and higher charged peptides between UnC and DADPS, UnC still had lower Xcorr scores, suggesting that the increase in highly charged states is not the primary cause of the difference in peptide identifications. Many of the detrimental characteristics of the AHA-UnC peptides we observed have been reported for TMT and other isobaric tags suggesting the large UnC tag is the major source of its poor performance in BONCAT analysis^28, 29, 37^. TMT label also appears to increase the difference between DADPS and UnC. This may be due to the presence of the large UnC modification and the large TMT tag on one peptide, which could further *reduce fragmentation efficiency* of the UnC peptides. We conclude that the improved performance of DADPS peptides in the DidBIT-BONCAT pipeline compared to UnC peptides is due to multifaceted effects.

## Acknowledgments

This work has been supported by NIH grants 5 R01 AG077046, 5R01AG075862 (to J.R.Y.), and P60 AA006420-41 We thank Dr. Claire Delahunty (TSRI) for critical review of the manuscript

## Author Contributions

Conceptualization, D.B.M and J.R.Y.; Investigation, D.B.M; Formal Analysis and Visualization, D.B.M; Writing – Original Draft, D.B.M.; Writing – Review & Editing, D.B.M.and J.R.Y.; Funding Acquisition and Project Administration, J.R.Y; Supervision, J.R.Y

## Supplementary Data

All MS raw files are uploaded on ProteomExchange PXD053959.

**Figure S1.**
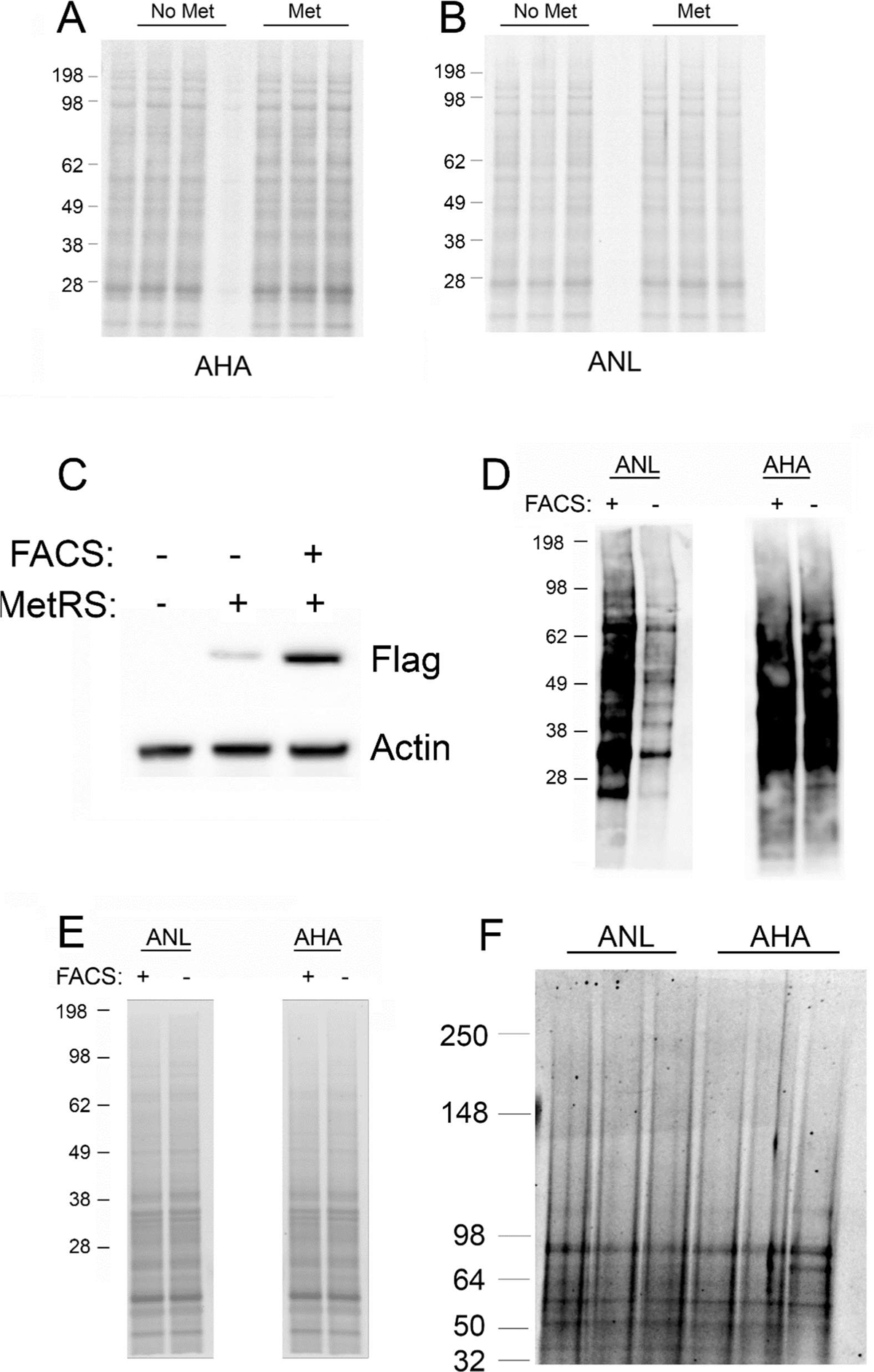
**A**, Total protein content of AHA samples in Fig1A using a protein stain. **B**, Total protein content of ANL samples in Fig1B using a protein stain. **C,** Immunoblot analysis of mMetRS transfected cells with and without FACS using a Flag tag antibody. **D**, Immunoblot analysis of ANL and AHA labeled cells with and without FACS probed with streptavidin-HRP. **E**, Total protein content of samples in D using a protein stain. **F**, Total protein content of samples in Fig. 1D using a protein stain.

**Figure S2.**
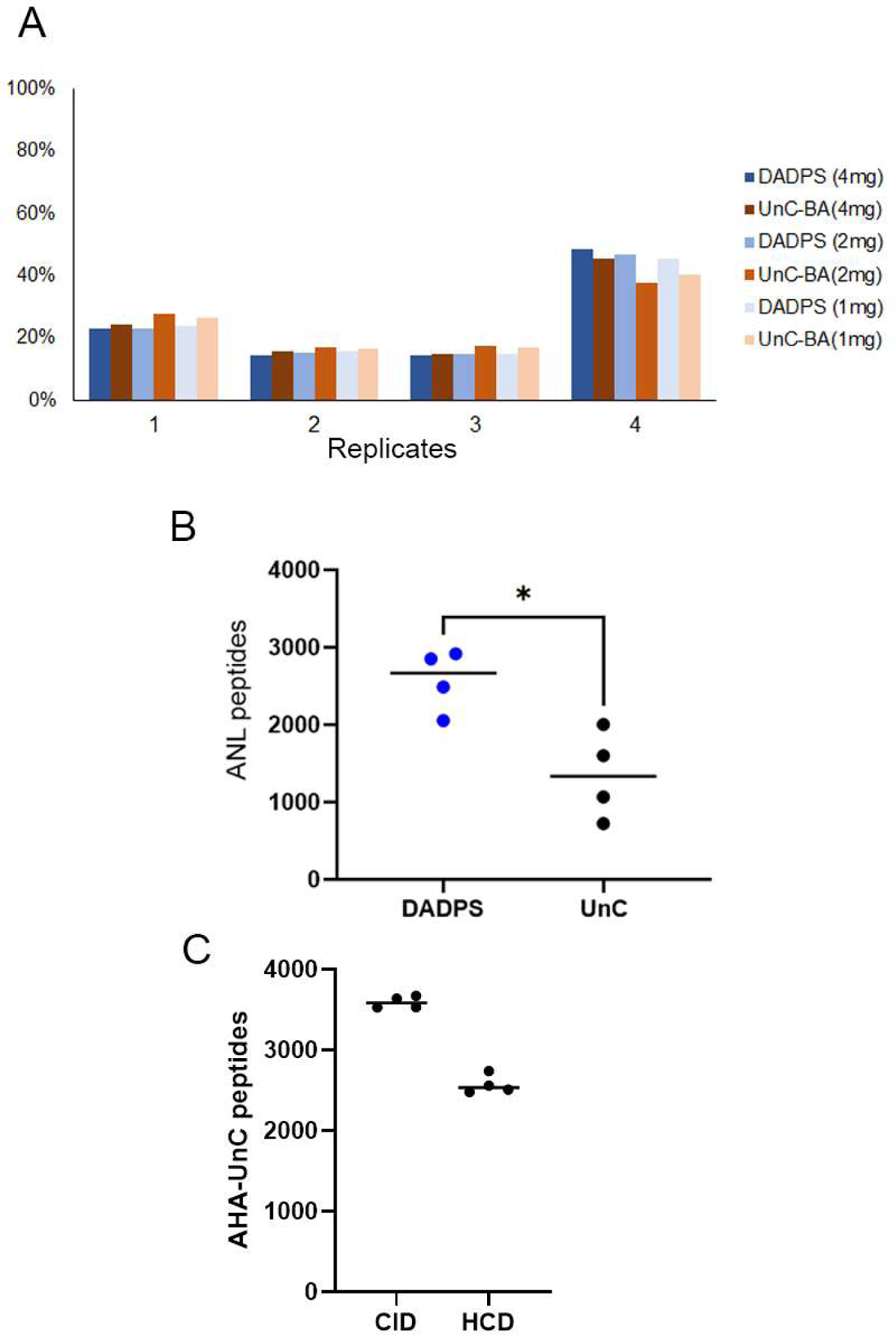
**A,** The number of replicates an AHA-BA peptide was identified by in Figure 2A. **B,** The number of ANL-BA peptides identification using either DADPS or UnC. **C,** One UnC-AHA sample was injected on an Orbitrap Fusion Lumos mass spectrometer (Thermo Fisher Scientific) using either an HCD or CID fragmentation in the ion trap.

